# Loss of PRICKLE1 in the myometrium leads to reduced fertility, abnormal myometrial architecture, and aberrant extracellular matrix deposition in mice

**DOI:** 10.1101/2024.09.12.612708

**Authors:** Emily R Roberts, Sornakala Ganeshkumar, Sumedha Gunewardena, Vargheese M Chennathukuzhi

**Affiliations:** Department of Cell Biology and Physiology, University of Kansas Medical Center, Kansas City, KS 66160, USA; Department of Biostatistics, University of Kansas Medical Center, Kansas City, KS 66160, USA

**Author notes:** Corresponding author: Vargheese M Chennathukuzhi. **Email:**. **Author Contributions:** V.M.C designed research; E.R.R., S. Ganeshkumar, and S. Gunewardena performed research; E.R.R., S. Gunewardena, and V.M.C. analyzed data; E.R.R. and V.M.C wrote and edited the manuscript. **Competing Interest Statement:** Authors declare no competing interest. **Classification:** Biological Sciences, Physiology.

**Keywords:** Prickle1, Wnt/PCP, uterine leiomyoma, fibroids, ECM

## Abstract

Uterine leiomyomas (UL) are the most prevalent benign tumors of the female reproductive tract, originating from the myometrium and affecting over 75% of reproductive-age women. Symptoms of UL include pelvic pain, pressure, dysmenorrhea, menorrhagia, anemia, and reproductive dysfunction. Currently, there is no effective long-term pharmacotherapy for UL, making them the leading cause of hysterectomies in the United States. The lack of treatment options is attributed to the absence of accurate animal models and a limited understanding of UL pathogenesis. Previous research has shown the loss of repressor of element 1 silencing transcription factor/neuron-restrictive silencing factor (REST/NRSF) within the myometrium promotes UL pathogenesis. In addition, deletion of *Rest* in the mouse myometrium leads to a UL phenotype. PRICKLE1, also known as Rest-interacting LIM-domain Protein (RILP), is required for nuclear localization of REST and Wnt/planar cell polarity (PCP) signaling, making it a critical target for UL studies. In the context of PCP, smooth muscle cells in UL show abnormal organization, aberrant ECM structure, and expression levels, potentially influenced by PRICKLE1 loss. The exact role of PRICKLE1 and Wnt/PCP in UL pathogenesis remains unclear.

To explore PRICKLE1’s role in UL, we deleted *Prickle1* using our myometrial-specific icre. Our findings demonstrate that *Prickle1* loss in the myometrium results in a UL phenotype characterized by altered collagen expression, excessive extracellular matrix (ECM) deposition, aberrant smooth muscle cell organization, increased *Esr1* and *Pgr* expression, and dysregulated Wnt/PCP signaling. This novel mouse model serves as a valuable preclinical tool for understanding UL pathogenesis and developing future pharmacotherapies.

## Introduction

Uterine leiomyomas (UL, uterine fibroids) are the most common benign tumors of the female reproductive tract, affecting over 75% of reproductive-age females (Bulun, 2013). Although benign, approximately 1 in 4 patients will have severe symptoms requiring clinical treatment (Bulun, 2013). Symptoms of UL include pelvic pain and pressure, dysmenorrhea, menorrhagia, anemia, and, in some instances, reproductive dysfunction (Al-Hendy and Salama, 2006, Cook et al., 2010). UL are hormone-responsive and arise from the myometrial smooth muscle cells of the uterus, ranging in size from 10 millimeters to over 20 centimeters (McWilliams and Chennathukuzhi, 2017).

Unfortunately, despite the large number of affected patients, there is no effective, long-term pharmacotherapy option to treat UL (Walker, 2002, Walker and Stewart, 2005). Due to this lack of options, UL has become the primary reason for hysterectomies, accounting for 39% of all hysterectomies performed in the United States (De La Cruz and Buchanan, 2017). The absence of long-term pharmacotherapies is primarily due to the lack of accurate animal models for UL and the gap in understanding the mechanisms behind UL pathogenesis. Previous work from our laboratory has demonstrated that the loss of NSRF/REST results in increased GPR10 and the activation of P13K/AKT-mTOR signaling in UL (Varghese et al., 2013). Moreover, ablation of *Rest* within the myometrium results in UL phenotype in mice with altered progesterone receptor function and enhanced estrogen signaling (Cloud et al., 2022).

PRICKLE1, also referred to as Rest-interacting LIM-domain Protein (RILP), is highly involved in nuclear localization of REST in addition to Wnt/planar cell polarity (PCP) signaling, making it an intriguing target of study in UL (Shimojo and Hersh, 2003, Katoh and Katoh, 2003). Previous work in the lab has shown the loss of PRICKLE1 in human UL samples compared to matched controls (McWilliams et al., 2024). Moreover, *in vitro* siRNA knockdown of *PRICKLE1* in human myometrial cells results in the loss of REST (McWilliams et al., 2024). In the context of planar cell polarity, UL structural proteins (collagens, fibronectins, and proteoglycans) are abnormal in organization, structure, and expression levels (Patel et al., 2016, Parker, 2007), potentially affected by the loss of PRICKLE1. The role PRICKLE1 plays within the myometrium in the context of UL is not well understood and may be key to further understanding of UL pathogenesis and the development of long-term pharmacotherapies.

To investigate the role of PRICKLE1 in UL, we developed a myometrial conditional knockout (cKO) of *Prickle1* using a myometrial specific icre (MiC) previously developed in our laboratory (Cloud et al., 2022). We demonstrate that the deletion of *Prickle1* in the myometrium leads to a UL phenotype consisting of fibrotic structures, excess extracellular matrix (ECM) production, altered ECM and smooth muscle cell organization, increased *Esr1* and *Pgr* expression, and dysregulated Wnt/PCP signaling. This novel mouse model provides a potentially crucial preclinical tool for investigating and understanding UL pathogenesis and the future development of pharmacotherapies.

## Materials and Methods

### Generation of myometrial-specific *Prickle1* conditional knockout mice (MiC mouse)

Mouse embryonic stem cell clone harboring floxed *Prickle1* (*Prickle1tm1a(EUCOMM)Wtsi,* ES cell clones with floxed exon 6 with conditional knockout potential) was acquired from EuCOMM. The ES cells were used to generate chimeric founder mice, which were mated to C57BL6 WT mice to create heterozygous *PRICKLE1*^fl/+^ mice. These mice were crossed with FLPo (Gt(ROSA)26Sor^tm2(FLP*)Sor^) mice to remove FRT flanked βgal-neo sequences. These were bred together to obtain homozygous *PRICKLE1^fl/fl^* mice, which were then crossed with transgenic myometrial-specific *iCre* (MiC) mice that we generated using the rat Calbindin D9K proximal promoter upstream of iCre, to produce *PRICKLE1* cKO mice, specific to the myometrium (Cloud et al., 2022). The mice were genotyped by PCR using primers for PRICKLE1 (5’GGTTTCATGTGTTGAGACATTTC) (5’GTATTTCTGTGCCCTTTTTGTCGTCG) (5’TGAACTGATGGCGAGCTCAGACC) as well as primers for MiC (5’CCACTAATGCTGTTCCGACCTGTC) (5’GGTGATGAGGAGAATCAGAAAGG) (5’CATCCTTGGCACCATAGATCAG). All animal procedures were performed per the Guide for Care and Use of Laboratory Animals, approved by the National Institute of Health (NIH), and followed an KUMC IACUC-approved animal study proposal.

### Histology and Staining

Uterine tissues were fixed in 4% paraformaldehyde for 24 hours and then processed for paraffin embedding. Tissues were deparaffinized in xylene and rehydrated in a series of ethanol washes and either stained for Hematoxylin and Eosin (H&E), immunofluorescence, or picrosirius red. For immunofluorescence staining, antigen retrieval was performed following rehydration by boiling in antigen unmasking solution (Vector Labs, Cat# H-3301). Following antigen retrieval, tissues were stained with primary antibodies anti-Col1a1 (Novus Biologicals, Cat# NB600-594SS, 1:100), anti-αSMA (Millepore, Cat# MABT381, 1:1000), anti-actin (Santa Cruz Biotechnology, Cat# sc-1616, 1:500), and DAPI. Alexa Fluor 488 secondary antibody was used for all immunofluorescence staining (Jackson Research Laboratories, Cat# 711-545-152, 1:200). Picrosirius red staining was performed via manufacture protocol within the kit (Polyscience Inc, Cat# 24901). All histological samples were imaged using a Nikon Eclipse 90i fluorescent microscope with Nikon Digital Sight color DS-Fi1 camera. Image analysis was performed using ImageJ to determine the corrected total cell fluorescence (CTCF) for 3 sections of 3 separate samples of 6-month-old cKO and controls in diestrus. Student’s nonparametric T-test was performed to determine statistical significance.

### RNA Isolation and qRT-PCR Analysis

Total RNA was isolated from uterine tissue samples stored in RNAlater (Invitrogen, Cat# AM7020) and then biopulvirized and placed into TRIzol Reagent (Invitrogen, Cat# 15596026). Following TRIzol, a series of chloroform, isopropanol, and ethanol washes were used to isolate the RNA. Following quantification using Nanodrop spectrophotometer, aliquots of RNA were reverse transcribed using High Capacity cDNA Reverse Transcription Kit (Applied Biosystems, 4368814). TaqMan assays for *Col1a1* (IDT, Mm.PT.47.9778198)*, Col4a1* (IDT, Mm.PT.58.31838522)*, Col4a2* (IDT, Mm.PT.58.28692624)*, Col6a3* (IDT, Mm.PT.58.15781337)*, Dpt* (IDT, Mm.PT.47.17098032)*, Acta2* (IDT, Mm.PT.47.7024949)*, Tgfb3* (IDT, Mm.PT.47.10648587), *Esr1* (IDT, Mm.PT.58.8025728), *Pgr* (IDT, Mm.PT.5810254276), *Prickle2* (IDT, Mm.PT.58.13127366), and *Prickle3* (IDT, Mm.PT.58.13792190) were used to quantify gene expression differences utilizing the delta delta C(T) method with housekeeping genes Rn18s (ThermoFisher Scientific, Mm03928990_G1) for 3 cKO and 3 control 6-month-old mice in diestrus.

### Fertility Assessment

Female *Prickle1^f/f^* MiC and *Prickle1^f/f^* mice were mated with *Prickle1^f/f^* males to induce pregnancy. The morning of finding the vaginal plug was considered day 0.5 of pregnancy. Litter size analysis was tracked via number of pups born following positive vaginal plug formation at days 17-21. Student’s nonparametric T-test was performed to determine statistical significance.

### Protein Extraction and Western Blotting

Frozen tissue samples were processed via biopulverization followed by sonication in Cell Lysis Buffer (Cell Signaling Technology, Cat# 9803) with protease and phosphatase inhibitor cocktails (ThermoFisher Scientific, Cat# P8340 and P0044). Samples were centrifuged at 18,000g at 4C for 15 minutes and the supernatant was collected. Protein was quantified using Bio-Rad Protein Assay Kit II (Bio-Rad, Cat# 50000002). 50 μg of protein was loaded with Laemmli buffer into 4-15% precast polyacrylamide gels (Bio-rad, Cat#5671083) and subjected to 130V for electrophoresis. Transfer onto a nitrocellulose membrane (ThermoFisher Scientific, Cat# 1620094) was performed at 4C for 1.5 hours at 70V. Membranes were blocked with 5% skim milk for 1 hour and incubated overnight at 4C with primary antibody. Primary antibodies included anti-mTOR (Cell Signaling Technology, Cat# 2983S, 1:1000), anti-AKT (Cell Signaling Technology, Cat# 9272S, 1:1000), anti-p-mTOR (Cell Signaling Technology, Cat# 2971S, 1:1000), anti-p-AKT (Cell Signaling Technology, Cat# 40605, 1:1000), anti-p-4EBP1 (Cell Signaling Technology, Cat# 94594, 1:1000), anti-p-P70S6 (Cell Signaling Technology, Cat# 42065, 1:1000), anti-LRP6 (Cell Signaling Technology, Cat# 3395S, 1:1000), anti-p-LRP6 (Cell Signaling Technology, Cat# 2568S, 1:500), anti-DVL1 (Sigma, Cat# D3570, 1:1000), anti-c-MYC (Cell Signaling Technology, Cat# 5605S, 1:1000), anti-DVL2 (Cell Signaling Technology, Cat# 3216S, 1:2000), anti-WNT3A (Cell Signaling Technology, Cat# C64F2, 1:1000), anti-SCRIBBLE (Cell Signaling Technology, Cat# 4475, 1:1000), anti-c-JUN (Cell Signaling Technology, Cat# 9165, 1:1000), anti-p-JNK (Santa Cruz Biotechnology, Cat# sc-6254, 1:1000), anti-p-PKC (Cell Signaling Technology, Cat# 9379S, 1:1000), anti-RAC1/2/3 (Cell Signaling Technology, Cat# 2465P, 1:1000), anti-RHOA (Cell Signaling Technology, Cat# 2117P, 1:1000), anti-WNT5A (Cell Signaling Technology, Cat# 2392, 1:1000), and anti-β-ACTIN (Genescript, Cat# A01865, 1:5,000). Following 3 washes, membranes were incubated at room temperature in anti-rabbit or anti-mouse IgG (Promega, Cat# W401B and W402B, 1:10,000) for 1 hour. Proteins were detected using SuperSignal West Pico PLUS Chemiluminescent Substrate (ThermoFisher Scientific, Cat# 34080). Blots were imaged using a BioRad Chemidoc MP imager. Western blot quantification was performed using ImageJ and normalized to loading controls for 3 cKO and 3 control 6-month-old mice in diestrus. Student’s nonparametric T-test was performed to determine statistical significance.

### Single-cell RNA Sequencing

Whole uteri of *Prickle1^f/f^* MiC (n=3) and *Prickle1^f/f^* (n=3) mice aged 6 months were isolated immediately following euthanasia using IACUCC approved methods and washed in cold 1X PBS. Uteri were digested as previously described (Cloud et al., 2022), and samples were pooled following digestion. The KUMC genomics core processed and sequenced the cells as previously described (Cloud et al., 2022). The *Prickle1^f/f^* MiC sample had 4998 cells with 38799 mean reads and 1431 median genes per cell, while the *Prickle1^f/f^* sample had 5426 cells with 43202 mean reads and 1098 median genes per cell. Both samples had high sequence saturation levels (> 60%) and the mapping rate of reads to the mouse genome (mm10) was > 90% for both samples. The data discussed in this publication have been deposited in NCBI’s Gene Expression Omnibus and are accessible through GEO Series accession number GSE276991.

### Single-cell Data Analysis

scRNA-sequencing libraries were generated using the 10x Chromium Single Cell 3’ v3 chemistry (10x Genomics) and sequenced in an Illumina NovaSeq 6000 sequencing machine. The raw sequence reads were processed using the 10x Genomics Cellranger pipeline (v 6.1.1) to obtain UMI feature-barcode count matrices. Doublets were removed using DoubletFinder (McGinnis et al., 2019) and cells filtered to contain only those cells with at least 500 UMIs and over 250 detected genes with a genes per UMI ratio greater than 0.8 and a mitochondrial gene ratio less than 20%. The resulting single-cell data was analyzed using the R software Seurat (v4), as previously described (Sasaki et al., 2024). The analysis was performed at a 0.4 cluster resolution reasoned using the Clustree software (Zappia and Oshlack, 2018), giving 14 stable clusters. The cluster cell types were identified using the SingleR software (Aran et al., 2019) and our expert curation as previously described (Cloud et al., 2022). Annotations were based on the two reference datasets, ImmGenData from the Immunological Genome Project (ImmGen) (Heng et al., 2008) and MouseRNAseqData (Benayoun et al., 2019).

### Ingenuity Pathway Analysis

Biological, functional, and pathway analysis were performed using Ingenuity Systems Pathway Analysis software (IPA, QIAGEN, Germantown, MD) on the significantly (FC >1.5, p-value <0.05) differentially expressed genes between *Prickle1^f/f^*MiC and *Prickle1^f/f^* myometrial (cluster 0), stromal (clusters 1 and 5), and epithelial (clusters 2 and 11) cells.

## Results

### cKO of *Prickle1* in Mouse Myometrium Leads to Uterine Leiomyoma Phenotype and Reduced Fertility

Using *Prickle1^f/+^* ES cells from EUCOMM (*Prickle1tm1a(EUCOMM)Wtsi)*, we developed *Prickle1^f/f^* mice and utilized myometrial-specific Cre (MiC) (Cloud et al., 2022) to conditionally ablate *Prickle1* in the mouse myometrium (Figure 1 A and B).

**Figure 1.**
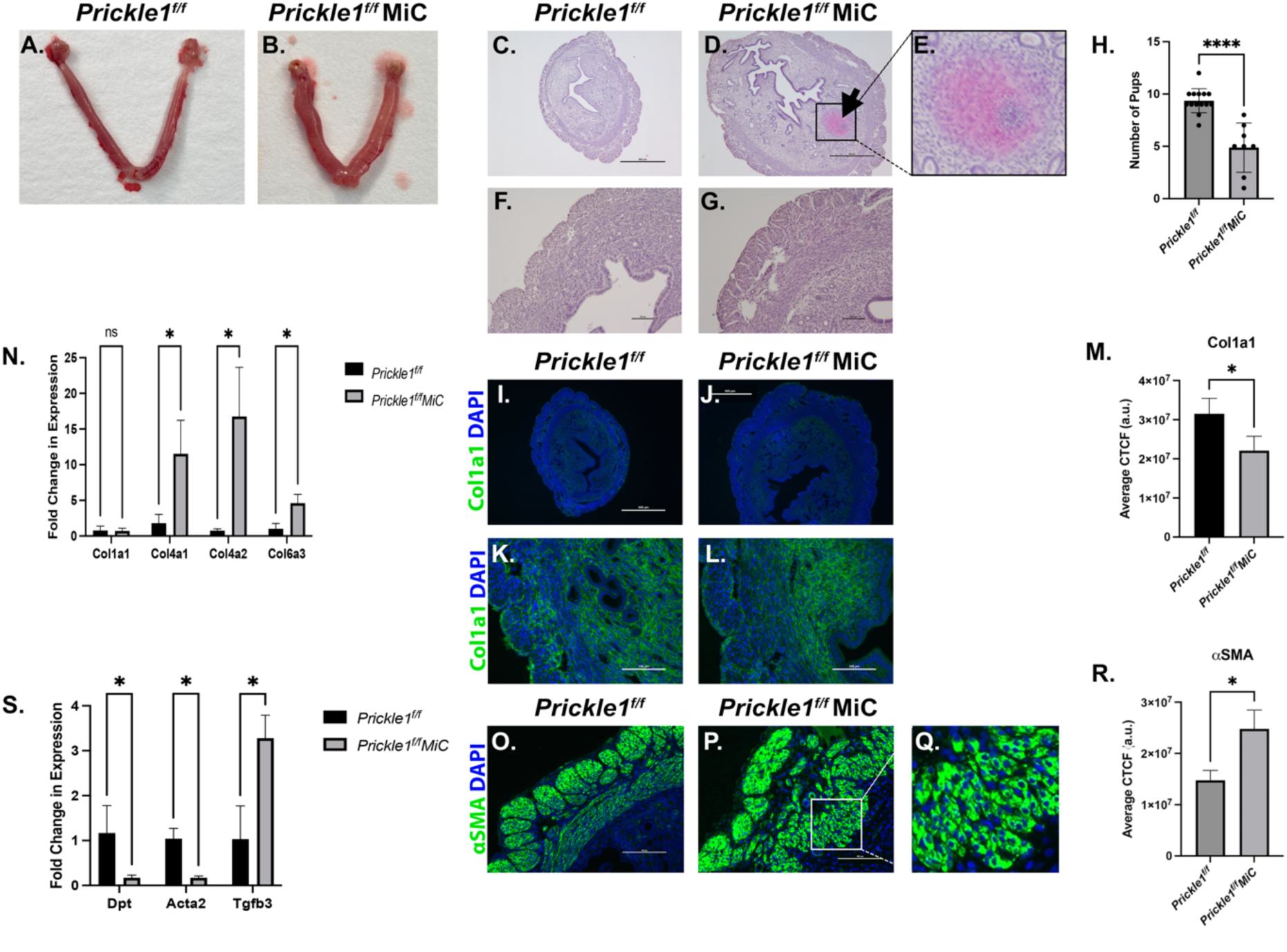
*Prickle1* cKO in mouse myometrium leads to Uterine Leiomyoma phenotype. (A-B) Representative image of (B) *Prickle1^f/f^* MiC mouse uteri compared to (A) control in diestrus. (C-G) H&E stain of uterine horn sections showing (C,F) *Prickle1^f/f^* and (D,E,G) *Prickle1^f/f^* MiC with fibrotic structure present in the (D) *Prickle1^f/f^* MiC as indicated by the arrow and (E) inset. (Scale bars: A and B 500 µm, C and D 100 µm). (H) Number of pups born to *Prickle1^f/f^* MiC (n=8) and control (n=23) female mice aged 6-months-old when bred with control males. Error bars represent ±SEM. Student’s nonparametric T-test was performed, ****P<0.0001. (I-L) Immunofluorescent images of the myometrium of control (I, K) and *Prickle1^f/f^* MiC (J,L) mice stained for Col1a1 (green) and DAPI (blue). (Scale bars: A and B 500 µm, C and D 100 µm). (M) Quantification of immunofluorescent images for Col1a1 showing decreased Col1a1 expression in the *Prickle1^f/f^* MiC cKO compared to control. Error bars represent ±SEM. Student’s nonparametric T-test was performed, *P<0.05. (N) TaqMan RT-qPCR analysis of collagen gene expression in uteri from 6-month-old *Prickle1^f/f^* MiC cKO compared to control. Genes include *Col1a1, Col4a1, Col4a2,* and *Col6a3*. Error bars represent ±SEM. Student’s nonparametric T-test was performed, *P<0.05. (O-Q) Immunofluorescent images of the myometrium of control (O) and *Prickle1^f/f^* MiC (P) mice stained for αSMA (green) and DAPI (blue) with inset (Q) showing the altered organization of circular smooth muscle cells in the cKO. (Scale bars, 100 µm). (R) Quantification of immunofluorescent images for αSMA showing increased expression in the *Prickle1^f/f^* MiC cKO compared to control. Error bars represent ±SEM. Student’s nonparametric T-test was performed, *P<0.05. (S) TaqMan RT-qPCR analysis in uteri from 6-month-old *Prickle1^f/f^* MiC cKO compared to control. Genes include *Dpt, Acta2, and Tgfb3.* Error bars represent ±SEM. Student’s nonparametric T-test was performed, *P<0.05.

Upon *Prickle1* ablation, we observed significant changes in the uterus, including altered tissue architecture in the myometrium and the formation of fibrotic structures within the endometrium (Figure 1 C-G). Furthermore, we noted a decrease in fertility in the cKO, consistent with a UL phenotype (Makker and Goel, 2013) (Figure 1 H). These changes were accompanied by alterations in ECM components, including a reduction in Col1a1 immunofluorescent staining (Figure 1 I-M) and an increase in *Col4a1*, *Col4a2*, and *Col6a3* RNA expression in the cKO, consistent with the UL phenotype of increased collagen deposition (Maekawa et al., 2013) (Figure 1 N). Additionally, immunofluorescent staining of alpha-smooth muscle actin (αSMA) showed increased expression in addition to circular myometrial smooth muscle cell altered organization into bundles in the cKO rather than a parallel organization, as seen in the control (Figure 1 O-Q).

Since UL is known to be associated with altered expression of genes involved in TGF-β signaling, decreased dermatoponin (*Dpt*), and altered alpha-smooth muscle actin (*Acta2*) (Arici and Sozen, 2000, Catherino et al., 2004, Wolanska et al., 1998) we performed TaqMan RT-qPCR to test the expression of *Dpt, Acta2,* and *Tgfb3.* Results from 6-month-old control and cKO mice indicated that cKO had significantly lower expression of *Dpt* and *Acta2* with increased expression of *Tgfb3*, consistent with UL phenotype (Figure 1 S). Western blot analysis of P13K-AKT/mTOR signaling pathway proteins indicated that this signaling cascade was not aberrantly activated in the cKO mice (Supplementary Figure 1), a phenotype distinct from that of the *Rest* cKO mouse model for UL (Cloud et al., 2022).

While alterations were seen from our analysis of the whole uterus, further understanding of individual cellular changes was pertinent. Therefore, to study the impact the loss of PRICKLE1 in the myometrium has on the whole uterus by cell type, single-cell RNA sequencing was performed on 6-month-old *Prickle1^f/f^*MiC and *Prickle1^f/f^* mice in diestrus (GSE276991).

The analysis identified 14 distinct clusters with differences in subpopulations between the control and the cKO (Figure 2 A). Analysis of collagen expression by cell type demonstrated increased *Col1a1* and *Col6a3* expression in stromal clusters (clusters 1 and 5) and epithelial clusters (clusters 2 and 11) in the cKO compared to control (Figure 2 B and C). Comparative u-MAP plots demonstrated increased expression of *Col3a1, Col4a1, Col4a2,* and *Col6a3* within the myometrial cluster (cluster 0) and stromal clusters (clusters 1 and 5) and increased *Col1a1* expression within stromal clusters in the cKO compared to control (Supplementary Figure 2). Further analysis demonstrated minimal alterations to REST target genes *Snap25, Gria2, Stmn2,* and *Mmp24* (Supplementary Figure 3). Whereas *Acta2* and *Tgfb3* expression were increased within the myometrial cluster (cluster 0) in the cKO compared to control (Supplementary Figure 3). In addition, increased expression of *Esr1* and *Pgr* was seen in both myometrial cluster 0 and stromal clusters 1 and 5, with increased *Esr1* expression seen in whole uterus samples, consistent with altered steroid hormone receptor expression seen in UL (Bulun, 2013, Borahay et al., 2015) (Figure 2 D-F).

**Figure 2.**
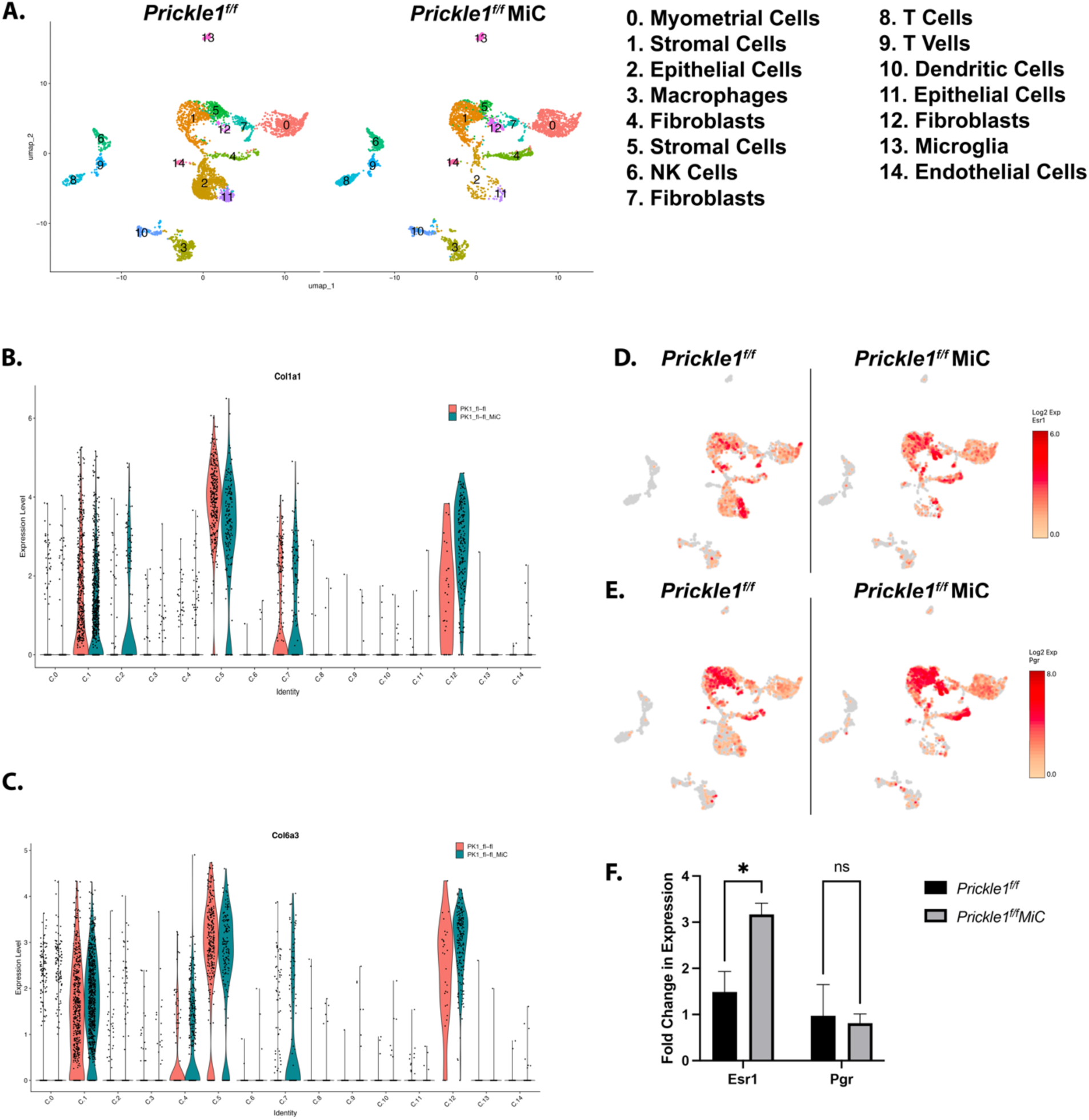
Single-cell RNA sequencing of *Prickle1* cKO in mouse myometrium demonstrates altered collagens, *Esr1*, and *Pgr* expression. (A) Individual u-MAP plots of the hierarchical clustering from uteri of 6-month-old controls and *Prickle1^f/f^* MiC cKO mice showing 14 distinct clusters. (B,C) Violin plots for (B) *Col1a1* and (C) *Col6a3* showing expression changes by cluster in control and *Prickle1^f/f^* MiC cKO. (D,E) Comparative u-MAP plots of control and *Prickle1^f/f^* MiC cKO from single-cell RNA sequencing showing expression (log 2-fold expression > 0) of (D) *Esr1* and (E) *Pgr.* (F) TaqMan RT-qPCR analysis of *Esr1* and *Pgr* expression in uteri from 6-month-old *Prickle1^f/f^* MiC cKO compared to control. Error bars represent ±SEM. Student’s nonparametric T-test was performed, *P<0.05.

Ingenuity pathway analysis (IPA) of the myometrial cluster (cluster 0) indicated upstream regulators to include several forms of vitamin D3 and ML290, an allosteric agonist of the relaxin receptor RXFP1 (Table 1). *Rxfp1* expression is higher in fibroblast clusters 4 and 12 of the cKO compared to control (Supplementary Figure 4). IPA also indicated dysregulation of genes involved in cellular functions of the myometrium (cluster 0), including collagen synthesis and degradation, endocrine system disorders, reproductive system disorders, skeletal and muscular disorders, and organ development (Tables 2, Supplementary Tables 1 and 2). IPA of stromal clusters 1 and 5 indicated dysregulation of genes involved in cell death and survival, molecular transport, organismal development and survival, development of carcinoma, advanced malignant tumor, and advanced stage tumor (Tables 3, Supplementary Tables 3-5). Moreover, IPA of epithelial clusters 2 and 11 displayed dysregulation of genes involved in pulmonary fibrosis, hepatic fibrosis, reproductive system disease, endocrine system disease, cancer, and cell growth and proliferation (Supplementary Tables 6 and 9).

**Table 1.** Pathway analysis of predicted upstream regulators associated with dysregulated genes in the myometrial cell populations of *Prickle1^f/f^* MiC cKO mice.

| Upstream Regulators | *P* value |
| --- | --- |
| 1alpha,20-dihydroxy vitamin D3 | 1.19E-06 |
| (20S,23)-dihydroxy vitamin D3 | 1.19E-06 |
| 1R,20,23-trihydroxy vitamin D3 | 1.19E-06 |
| 20S-hydroxyvitamin-D3 | 1.19E-06 |
| ML290 | 2.45E-06 |

Pathway predictions made by IPA software based off genes which were found to be dysregulated in myometrial cells of 6-month-old *Prickle1<sup>ff</sup>* *MiCre* mice cKO (n=3) in diestrus.

### cKO of *Prickle1* in Mouse Myometrium Leads to Dysregulation of Wnt signaling

Due to PRICKLE1’s involvement in non-canonical Wnt/planar cell polarity (PCP) signaling, we performed a western blot analysis of whole uterus samples for both canonical and non-canonical Wnt signaling (Figure 3). This analysis indicated that canonical Wnt signaling was not activated (Figure 3 A and B) even though the canonical Wnt signaling pathway activation is still highly debated in UL. The non-canonical Wnt/PCP signaling analysis showed that the pathway was not significantly affected by the loss of PRICKLE1, except for lower SCRIBBLE expression and increased PKC phosphorylation in the cKO compared to the control (Figure 3 C-M). This may be due to species-specific compensation from Prickle2 and Prickle3, as evidenced by increased RNA expression within the cKO compared to control (Supplementary Figure 5).

**Figure 3.**
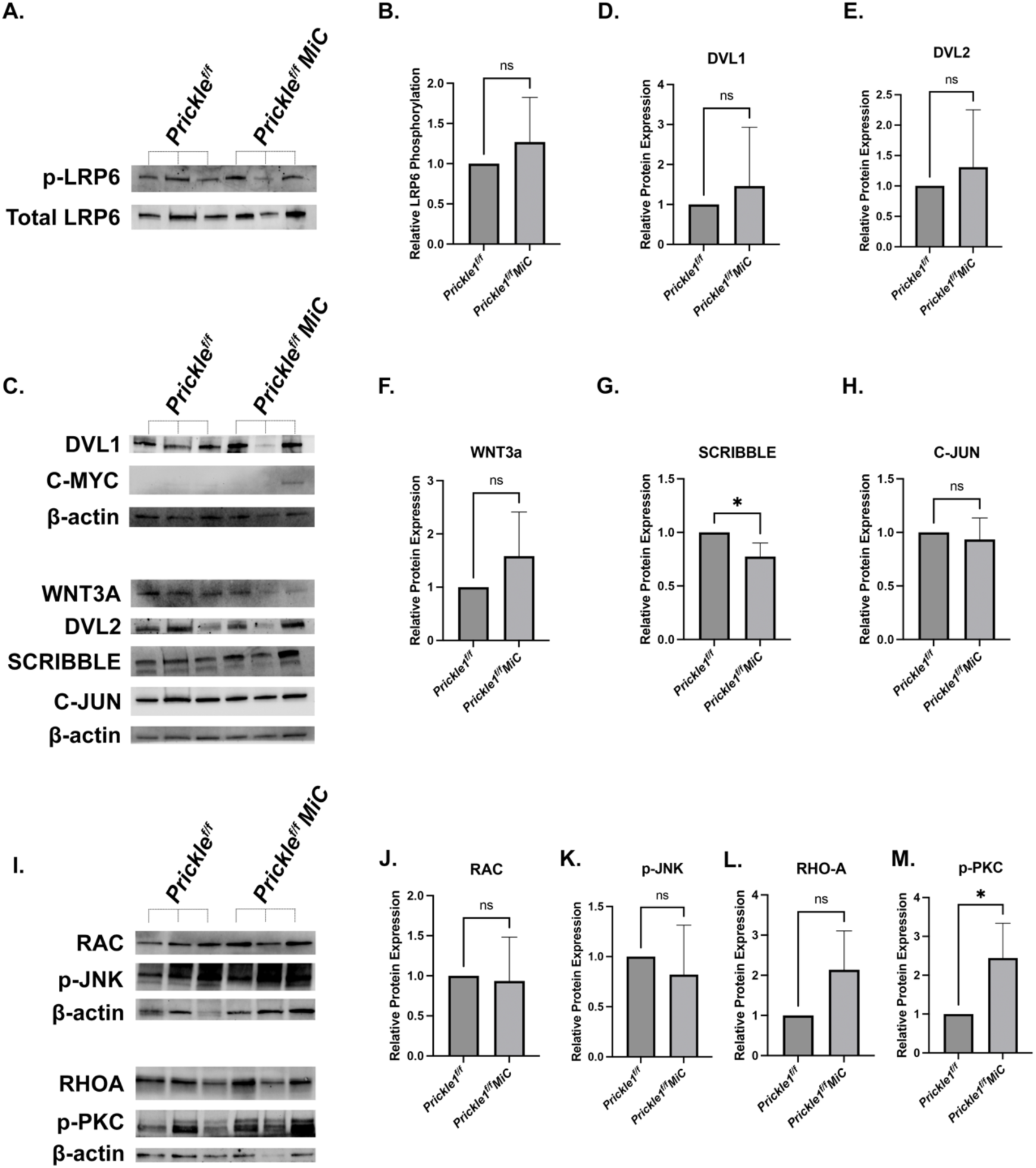
Wnt signaling pathway assessment in *Prickle1^f/f^* MiC conditional knockout. (A) Western blot showing canonical Wnt signaling phosphorylation of LRP6 with total LRP6 as a loading control. (B) Densitometric analysis of Western blot for phosphorylation of LRP6 with total LRP6 as a loading control. (C) Western blot showing non-canonical Wnt signaling pathway proteins with β-actin as a loading control. (D-H) Densitometric analysis of Western blot for non-canonical Wnt signaling pathway with β-actin as a loading control. (I) Western blot showing additional direct targets of non-canonical Wnt signaling pathway proteins with β-actin as a loading control. (J-M) Densitometric analysis of Western blot for direct targets of non-canonical Wnt signaling pathway with β-actin as a loading control. Error bars represent ±SEM. Student’s nonparametric T-test was performed, *P<0.05.

However, when looking at specific cell types, there is strong evidence for dysregulation in Wnt signaling pathways. Comparative u-MAP plots from the single-cell RNA sequencing analysis indicated increased *Dvl1*, *Dvl2, Scrib, Nkd1,* and *Nkd2* expression in stromal clusters 1 and 5 with decreased expression within the epithelial clusters (clusters 2 and 11) in the cKO compared to control (Figure 4 A-E). Moreover, *Vangl1, Vangl2,* and *Wnt5a* expression were minimally altered with small decreases in epithelial clusters (clusters 2 and 11) and slight increases in stromal clusters (clusters 1 and 5) in the cKO compared to control (Figure 4 F-H).

**Figure 4.**
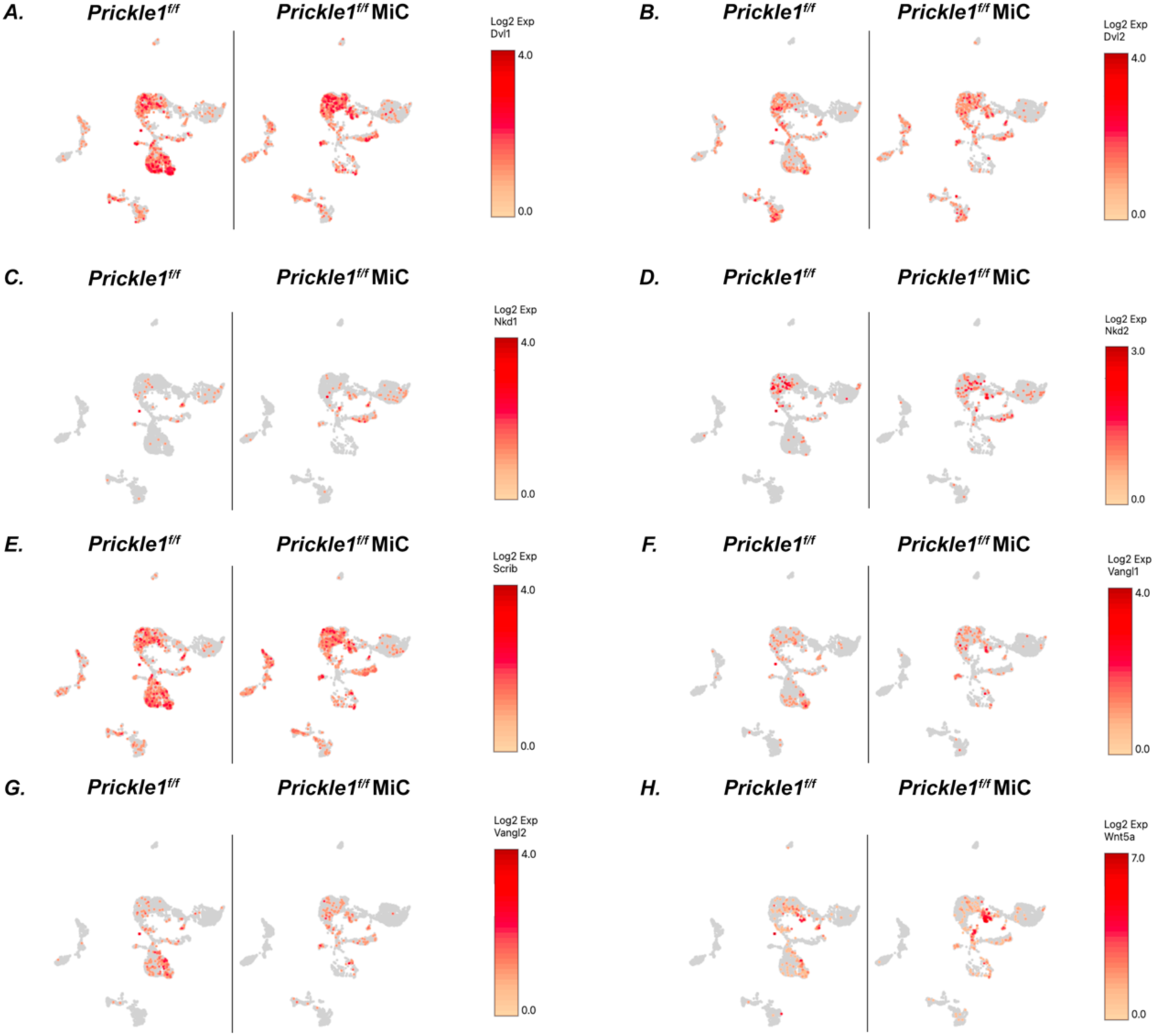
Single-cell RNA sequencing analysis of Wnt signaling genes. Comparative u-MAP plots of control and *Prickle1^f/f^* MiC cKO from single-cell RNA sequencing showing expression (log 2-fold expression > 0) of Wnt signaling genes (A) *Dvl1,* (B) *Dvl2,* (C) *Nkd1,* (D) *Nkd2,* (E) *Scrib,* (F) *Vangl1,* (G) *Vangl2,* and (H) *Wnt5a*.

### cKO of *Prickle1* in Mouse Myometrium Leads to Dysregulation of Cellular Organization

In addition to the altered cellular organization of the circular smooth muscle cells of the myometrium into bundles seen via αSMA staining (Figure 1 N and O), immunofluorescence staining for total actin displayed decreased levels of actin in addition to a similar alteration in organization in the cKO (Figure 5 A-E) Additionally, Picrosirius Red staining indicated alterations to collagen organization and increased collagen deposition levels in the cKO compared to the control (Figure 5 F-I), a hallmark of UL. The cKO showed collections of bundles of collagen fibers throughout the myometrium and stroma rather than parallel and longitudinal fibers, as seen within the control.

**Figure 5.**
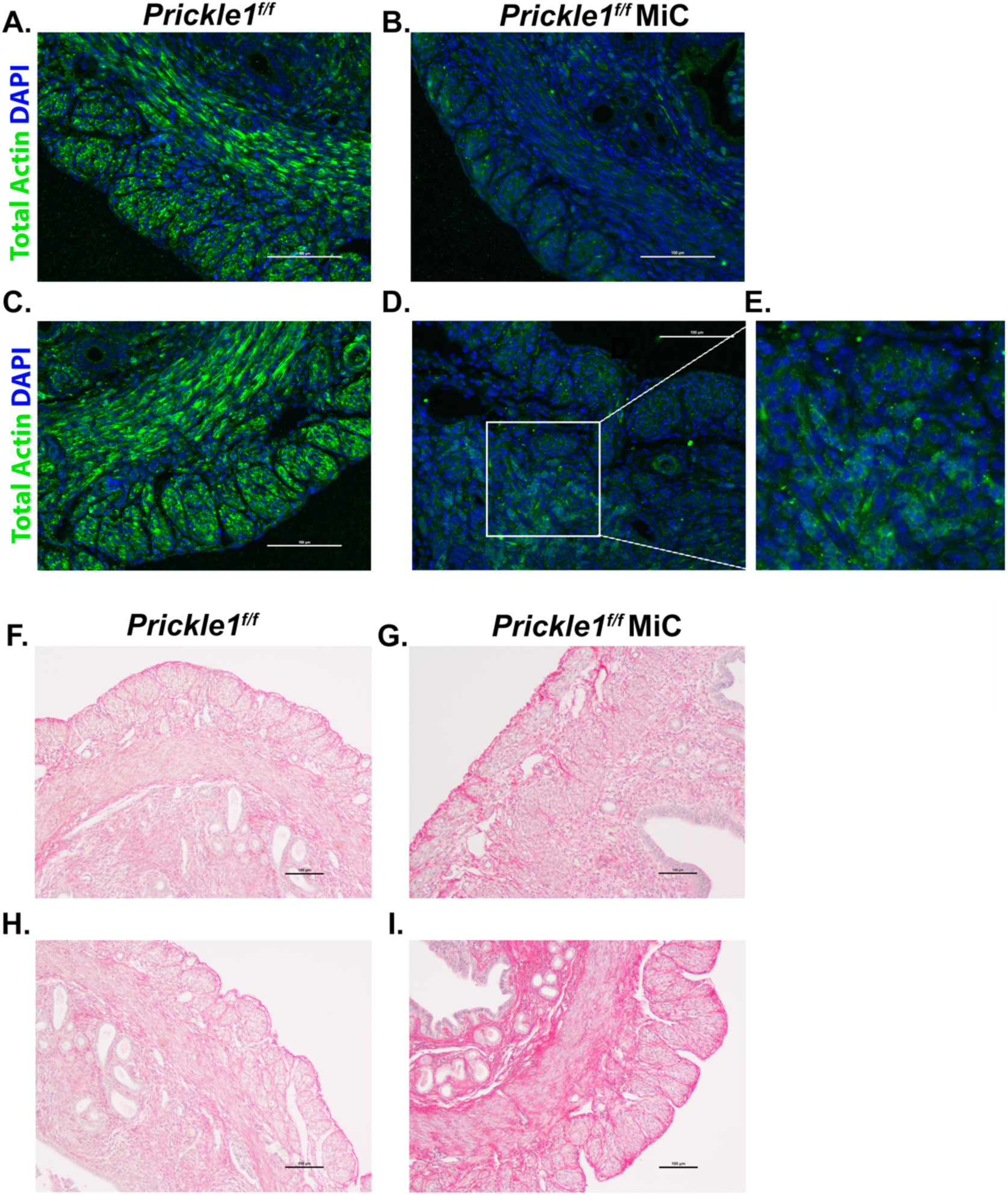
*Prickle1* cKO in mouse myometrium leads to dysregulation of cellular and ECM organization. (A-D) Immunofluorescent images of the myometrium of control (A,C) and *Prickle1^f/f^* MiC (B, D) mice stained for total actin (green) and DAPI (blue). (Scale bars: A and B 500 µm, C and D 100 µm). E. Inset of *Prickle1^f/f^* MiC mice stained for total actin (green) and DAPI (blue) showing altered organization of circular smooth muscle cells. (F-I) Picrosirius Red staining of the myometrium of control (F.H) and *Prickle1^f/f^* MiC (G,I) mice. (Scale bars 100 µm).

In addition, IPA of the myometrium (cluster 0) included dysregulation of collagen chain trimerization, assembly of collagen fibrils, ECM organization, and tissue development (Table 2, Supplementary Tables 1 and 2). IPA of stromal clusters 1 and 5 indicated dysregulation of genes involved in cellular movement, cell death and survival, organismal development, connective tissue development and function, and tissue morphology (Supplementary Tables 3-5). Moreover, IPA of epithelial clusters 2 and 11 displayed dysregulation of genes involved in ECM organization, integrin cell surface interactions, assembly of collagen fibrils, cellular movement, cell death and survival, cellular development, cellular growth and proliferation, cell-to-cell signaling, and tissue morphology (Supplementary Tables 6 and 9).

**Table 2.** Pathway analysis of predicted top canonical pathways associated with dysregulated genes in the myometrial cell population (cluster 0) of *Prickle1^f/f^* MiC cKO mice.

| **Canonical Pathways** | ***P* value** |
| --- | --- |
| Collagen biosynthesis and modifying enzymes | 6.42E-07 |
| Collagen chain trimerization | 1.24E-05 |
| Collagen degradation | 3.34E-05 |
| Assembly of collagen fibrils and other multimeric structures | 3.34E-05 |
| Extracellular matrix organization | 1.74E-04 |
| Pathway predictions made by IPA software based off genes which were found to be dysregulated in myometrial cells of 6-month-old *Prickle1<sup>ff</sup>* *MiCre* mice cKO (n=3) in diestrus. |  |

## Discussion

A better understanding of the mechanisms affecting the pathogenesis of UL is critical to developing pharmacotherapies for treating UL. There is a vast need for better animal models of UL that encompass the various characteristics of UL. Previously, our lab has shown that the loss of REST in the myometrium leads to increased expression of GPR10 and activation of the PI3K/AKT-mTOR pathway (Varghese et al., 2013). Furthermore, we have previously demonstrated that ablation of *Rest* in the myometrium results in UL phenotype in mice (Cloud et al., 2022). However, these current models require more information regarding the mechanisms behind the loss of REST and the cellular and ECM organizational alterations characteristic of UL.

PRICKLE1, also known as Rest-interacting LIM-domain Protein (RILP), has been previously shown to facilitate nuclear localization of REST (Shimojo and Hersh, 2003). Previously, we showed that PRICKLE1 is lost in UL and corresponds with the loss of REST (McWilliams et al., 2024). Moreover, PRICKLE1 is a crucial component of the non-canonical Wnt/Planar Cell Polarity (PCP) signaling cascade (Adler, 2012, Butler and Wallingford, 2017). PCP proteins, and particularly PRICKLE1, have been understudied in the context of UL.

Here, we developed a novel *Prickle1* myometrial-specific conditional knockout mouse model that, at least in part, mimics UL phenotype and provides a better understanding of the role PCP plays in UL pathogenesis and uterine physiology. Our *Prickle1^f/f^* MiC cKO mice showed fibrotic structures within the endometrium at 6 months old (Figure 1 D and E). Further investigation of these structures is necessary to determine their morphogenesis. However, these structures are not seen in the controls. Furthermore, alterations to collagen deposition and changes to ECM composition are characteristic of UL phenotype (Walker and Stewart, 2005, Maekawa et al., 2013). Our cKO showed a reduction in COL1A1 protein expression (Figure 1 I-M) with an increase in *Col3a1, Col4a1, Col4a2,* and *Col6a3* RNA expression (Figure 1 N, Figure 2 B and C, and Supplementary Figure 2), consistent with this UL phenotype. Reduced *Col1a1* in the ckO may be due to species-specific differences between rodent and human. Additionally, alteration to αSMA organization and expression in the ckO (Figure 1 O-Q) is consistent with UL phenotypic behavior (Kobayashi et al., 1996, Wolanska et al., 1998). Differences seen in the *Acta2* RNA, with decreased expression in the cKO, is likely due to the use of whole uterus samples for RNA analysis. Moreover, increased expression of TGF-β3 through decreased *Dpt*, a known regulator of TGF-β3, is an established component of UL (Arici and Sozen, 2000, Kuroda et al., 1999). Our cKO model shows both a decrease in *Dpt* with an associated increase in *Tgfb3*, consistent with this phenotype (Figure 1 S). Increased expression of *Esr1* and *Pgr* in the cKO (Figure 2 D-F) is also consistent with a UL phenotype (Borahay et al., 2015, Bulun, 2013), allowing for the improved study of hormone sensitivity and altered response to steroid hormones seen in UL. Previous work within our lab has shown similar increases in *Esr1* and *Pgr* in REST cKO mouse models (Cloud et al., 2022), although an epistatic analysis of this dysregulation needs further research.

In addition, IPA of differentially expressed genes within myometrial, stromal, and epithelial clusters indicated upstream regulators, including various forms of vitamin D3 and ML290. Vitamin D3 deficiency is a well-established risk factor for UL, and vitamin D3 has previously been implicated for possible use as a therapeutic for UL (Baird et al., 2013, Sabry et al., 2013). This provides another potential connection to known pathways in UL involving vitamin D3 and the loss of PRICKLE1. ML290 is an allosteric agonist of the relaxin receptor RXFP1 and exhibited anti-fibrotic properties in cardiac fibroblasts (Kocan et al., 2017), making a possible connection for its regulation and, coupled with the Rxfp1 expression in the cKO (Supplementary Figure 4), its potential therapeutic use in UL. Furthermore, IPA showed dysregulation of pathways associated with several cellular processes, fibrosis, and reproductive tumors and disorders, providing even further connection to the loss of PRICKLE1 and UL pathogenesis (Tables 1-3, Supplementary Tables 1-9). Dysregulated pathways that include pulmonary fibrosis and hepatic fibrosis (Supplementary Table 6) are highly associated with and indicative of the fibrosis seen in UL, even though these changes are seen within the epithelial cell population of the cKO. Pathway dysregulation of cell death and survival, carcinoma, and tumor formation are potentially related to UL tumorigenesis (Supplementary Tables 3 and 5). The majority of the changes indicated with IPA were seen within the stromal clusters, consistent with stromal changes seen in UL that lead to symptoms of excessive bleeding and increased inflammation (Zheng et al., 2014).

**Table 3.** Pathway analysis of predicted top diseases and disorders associated with dysregulated genes in the stromal cell population (clusters 1 and 5) of *Prickle1^f/f^* MiC cKO mice.

| **Diseases and Disorders** | ***P* value** |
| --- | --- |
| Metabolic disease | 9.83E-11 |
| Organismal injury and abnormalities | 9.83E-11 |
| Cancer | 1.01E-09 |
| Reproductive system disease | 3.17E-09 |
| Endocrine system disorders | 1.14E-08 |

Disease and disorder predictions made by IPA software based off genes which were found to be dysregulated in stromal cells of 6-month-old *Prickle1<sup>ff</sup>* *MiCre* mice cKO (n=3) in diestrus.

It is known that PI3K-AKT/mTOR signaling is activated in UL (Crabtree et al., 2009) and is increased in the REST cKO UL mouse model (Cloud et al., 2022). However, our cKO did not demonstrate activation of PI3K-AKT/mTOR signaling (Supplementary Figure 1). We also did not see any significant alterations to REST target genes as expected (Supplementary Figure 3). This may be due to rodent-specific differences in how PRICKLE1 regulates REST and requires further investigation. Furthermore, it is unknown if the increased *Prickle2* and *Prickle3* expression in the cKO model could compensate for REST stabilization.

In the context of Wnt/PCP signaling, the loss of PRICKLE1 in our cKO showed alterations to non-canonical Wnt signaling pathways at both the protein (Figure 3) and RNA (Figure 4) levels with subsequent changes to myometrial and ECM organization (Figures 1 and 5). Small changes in Wnt signaling protein expression is likely due to using whole uterus samples for evaluation with compensation from other Prickle homologs (Supplementary Figure 5). Single-cell RNA sequencing demonstrates alteration to RNA expression in myometrial clusters for several Wnt signaling genes with corresponding compensation in stromal clusters, likely preventing visualization of protein changes within whole uterus samples. Previously, *in vitro* studies have demonstrated structural changes occur in primary myometrial cells to allow for parallel organization, while leiomyoma cells become aggregates and form bundled nodules (Kobayashi et al., 1996). Our cKO mimics this behavior through alterations in ECM and myometrial cell organization, likely due to the disruption of Wnt/PCP signaling through the ablation of *Prickle1*. Altered architecture seen in the cKO would also indicate that potential remodeling of Wnt/PCP medicated cytoskeleton organization and tissue architecture changes occur even after puberty when *iCre* is expressed in MiC (Cloud et al., 2022). Alterations to myometrial and ECM organization, coupled with alterations to ECM within the stroma, may play a role in the fertility phenotype seen within the cKO (Figure 1 G) and in particular cases of UL (Al-Hendy and Salama, 2006). Moreover, IPA analysis of myometrial, stromal, and epithelial differentially expressed genes provides further evidence for Wnt/PCP disruption through the dysregulation of pathways involved in cell communication, tissue organization, ECM organization, collagen fibril organization, connective tissue morphology, and protein expression in myometrial, stromal, and epithelial cell populations (Tables 1-3, Supplementary Tables 1-9).

Our novel *Prickle1^f/f^* MiC cKO links the loss of PRICKLE1 to UL pathogenesis by disrupting Wnt/PCP signaling, collagen expression, and myometrial ECM organization. Our cKO developed fibrotic structures and contains many of the same human UL characteristics, making it an exciting model for the study of UL pathogenesis. Our model allows for further research of Wnt/PCP cellular communication and organization to the subsequent pathways in UL, providing an important pre-clinical model to study UL pathogenesis. Further analysis of this mouse model and the single-cell RNA sequencing data will be crucial to better the field’s understanding of the relationship between UL, PRICKLE1, and Wnt/PCP signaling, ultimately leading to better development of pharmacotherapies for the treatment of UL. This study further establishes that the role of Wnt/PCP dysregulation, rather than the canonical Wnt signaling pathway, is important for the development of UL.

## Acknowledgments and Funding Sources

V.M.C. was supported by grants from the NIH: P20 RR016475, R01 HD094373, R01 HD076450, R01 HD105714-01. E.R.R. was supported by the Madison and Lila Self Graduate Fellowship.

The authors acknowledge the Transgenic and Gene-Targeting Institutional Facility for help with the transgenic mice, the Genomics Core (Kansas Intellectual and Developmental Disability Research Center (KIDDRC) (NIH U54 HD090216), the Centers of Biomedical Research Excellence (P30 GM122731-03), and the NIH S10 High-End Instrumentation Grant (NIH grants S10 OD021743)) for help with the scRNA sequencing, and the KIDDRC (NIH U54 HD 090216) for help with the imaging at the University of Kansas Medical Center, Kansas City, KS, 66160.

