## Supplemental Information for "Loss of PRICKLE1 in the myometrium leads to reduced fertility, abnormal myometrial architecture, and aberrant extracellular matrix deposition in mice"

*Corresponding author: Vargheese M Chennathukuzhi.

**This PDF file includes:**

Figures S1 to S4
Tables S1 to S9

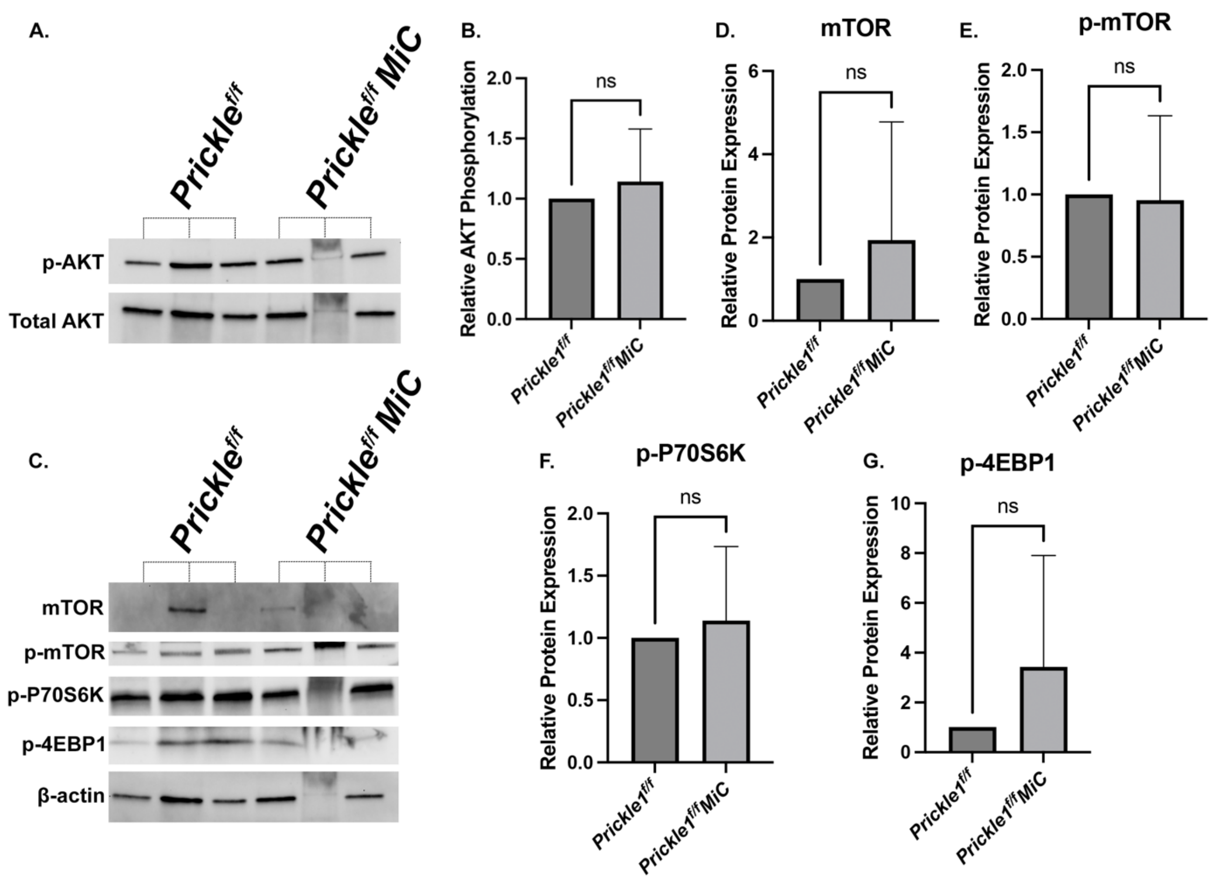


**Supplemental Figure 1. PI3K-AKT/mTOR pathway assessment in *Prickle1^f/f^* MiC conditional knockout.** (A) Western blot showing phosphorylation of AKT (s473) with total AKT as a loading control. (B) Densitometric analysis of Western blot for phosphorylation of AKT with total AKT as a loading control. (C) Western blot showing mTOR signaling pathway proteins with β-actin as a loading control. (D-G) Densitometric analysis of Western blot for mTOR signaling pathway with β-actin as a loading control. Error bars represent ±SEM. Student's nonparametric T-test was performed.

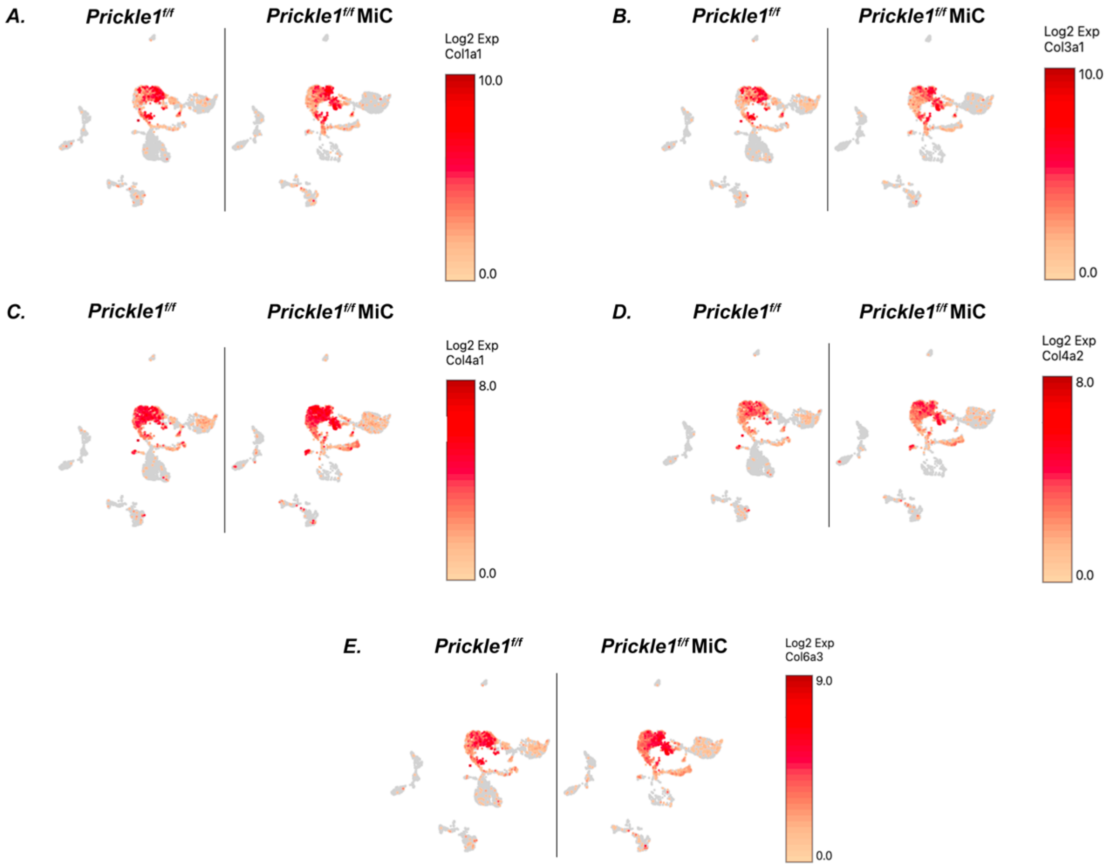


**Supplemental Figure 2. Single-cell RNA sequencing analysis of collagens.** Comparative u-MAP plots of control and *Prickle1$^{f/f}$* MiC cKO from single-cell RNA sequencing showing expression (log 2-fold expression > 0) of collagens including (A) *Col1a1,* (B) *Col3a1,* (C) *Col4a1,* (D) *Col4a2,* and (E) *Col6a3.*

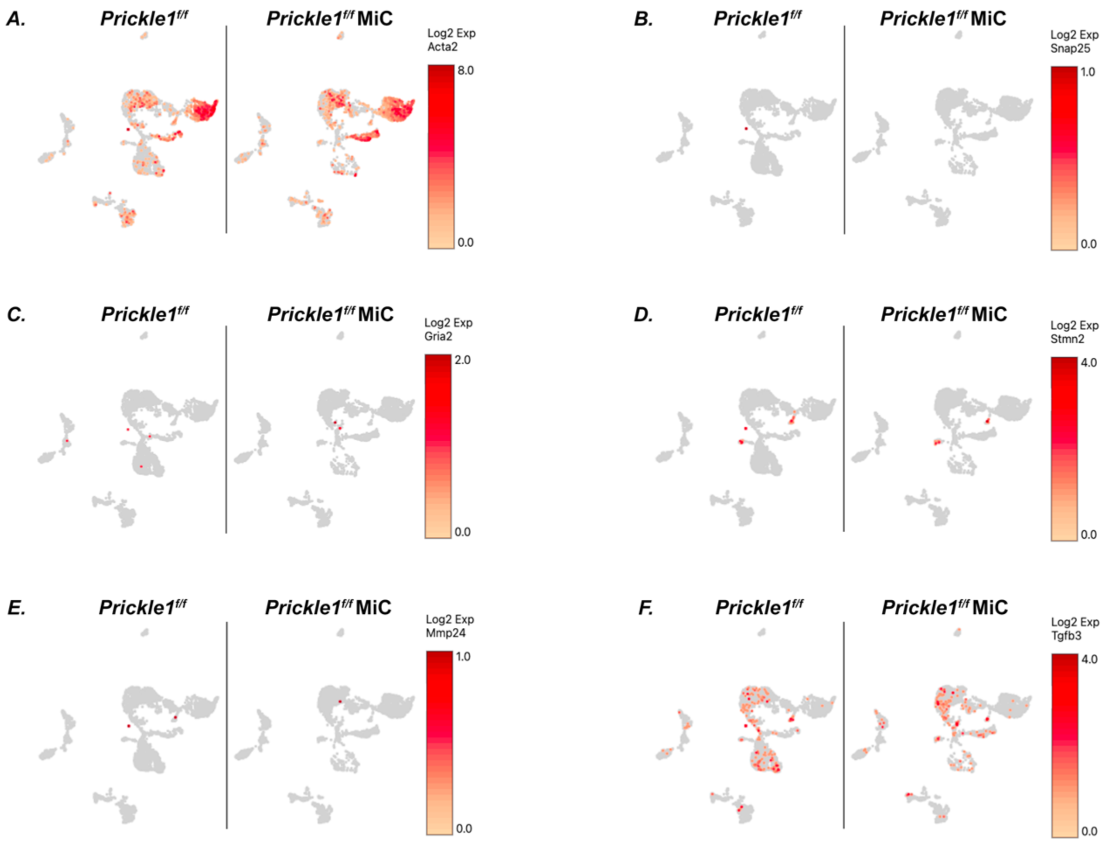


**Supplemental Figure 3. Single-cell RNA sequencing analysis of REST target genes.**

Comparative u-MAP plots of control and *Prickle1*$^{f/f}$ MiC cKO from single-cell RNA sequencing showing expression (log 2-fold expression > 0) of REST target genes (A) *Acta2,* (B) *Snap25,* (C) *Gria2,* (D) *Stmn2,* (E) *Mmp24,* and (F) *Tgfb3*.

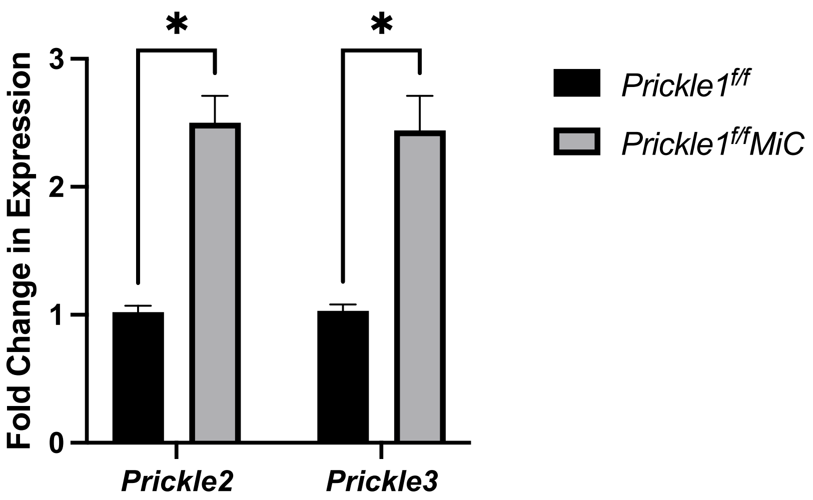


**Supplemental Figure 4. Gene expression of *Prickle2* and *Prickle3*** in 6-month-old *Prickle1*$^{f/f}$ MiC mice compared with control (n=3). Error bars represent ± SEM. Student's *t* test was performed, *$P<0.05$.

**Supplemental Table 1. Pathway analysis of predicted diseases and disorders associated with dysregulated genes in the myometrial cell population (cluster 0) of *Prickle1*$^{f/f}$ MiC cKO mice.**

| **Diseases and Disorders** | ***P* value** |
|---|---|
| Cancer | 2.31E-07 |
| Endocrine system disorders | 2.31E-07 |
| Organismal injury and abnormalities | 2.31E-07 |
| Reproductive system disease | 2.31E-07 |
| Skeletal and muscular disorders | 3.09E-07 |

Function predictions made by IPA software based off genes which were found to be dysregulated in myometrial cells of 6-month-old *Prickle1*$^{f/f}$ *MiCre* mice cKO (n=3) in diestrus.

**Supplemental Table 2. Pathway analysis of predicted physiological system development and functions associated with dysregulated genes in the myometrial cell population (cluster 0) of *Prickle1*$^{f/f}$ MiC cKO mice.**

| Physiological System Development and Function | *P* value |
|---|---|
| Tissue development | 1.64E-04 |
| Embryonic development | 1.96E-04 |
| Organismal development | 1.96E-04 |
| Organismal functions | 2.15E-04 |
| Organ development | 2.32E-04 |

Function predictions made by IPA software based off genes which were found to be dysregulated in myometrial cells of 6-month-old *Prickle1*$^{f/f}$ *MiCre* mice cKO (n=3) in diestrus.

**Supplemental Table 3. Pathway analysis of predicted molecular and cellular functions associated with dysregulated genes in the stromal cell population (clusters 1 and 5) of *Prickle1*$^{f/f}$ MiC cKO mice.**

| **Molecular and Cellular Functions** | ***P* value** |
|---|---|
| Cellular movement | 4.68E-11 |
| Carbohydrate metabolism | 4.44E-10 |
| Cell death and survival | 2.70E-09 |
| Molecular transport | 5.15E-08 |
| Small molecule biochemistry | 5.15E-08 |

Function predictions made by IPA software based off genes which were found to be dysregulated in stromal cells of 6-month-old *Prickle1*$^{f/f}$ *MiCre* mice cKO (n=3) in diestrus.

**Supplemental Table 4. Pathway analysis of predicted physiological system development and functions associated with dysregulated genes in the stromal cell population (clusters 1 and 5) of *Prickle1*$^{f/f}$ MiC cKO mice.**

| **Physiological System Development and Function** | ***P* value** |
|---|---|
| Cardiovascular system development and function | 8.91E-11 |
| Organismal development | 8.91E-11 |
| Organismal survival | 6.32E-09 |
| Connective tissue development and function | 3.96E-08 |
| Tissue morphology | 3.96E-08 |

Function predictions made by IPA software based off genes which were found to be dysregulated in stromal cells of 6-month-old *Prickle1*$^{f/f}$ *MiCre* mice cKO (n=3) in diestrus.

**Supplemental Table 5. Pathway analysis of top ML disease pathways associated with dysregulated genes in the stromal cell population (clusters 1 and 5) of *Prickle1$^{f/f}$* MiC cKO mice.**

| **Top ML Disease Pathways** | ***P* value** |
|---|---|
| Angiodema | 1.09E-04 |
| Development of carcinoma | 1.70E-04 |
| Advanced malignant tumor | 2.08E-04 |
| Advanced stage tumor | 2.08E-04 |
| Glomerulosclerosis | 2.29E-04 |

Pathway predictions made by IPA software based off genes which were found to be dysregulated in stromal cells of 6-month-old *Prickle1$^{f/f}$ MiCre* mice cKO (n=3) in diestrus.

**Supplemental Table 6. Pathway analysis of top canonical pathways associated with dysregulated genes in the epithelial cell population (clusters 2 and 11) of *Prickle1$^{f/f}$* MiC cKO mice.**

| **Top Canonical Pathways** | ***P* value** |
|---|---|
| Extracellular matrix organization | 2.51E-22 |
| Pulmonary fibrosis idiopathic signaling pathways | 1.75E-15 |
| Hepatic fibrosis/hepatic stellate cell activation | 4.43E-15 |
| Integrin cell surface interactions | 9.39E-15 |
| Assembly of collagen fibrils and other multimeric structures | 8.08E-13 |

Pathway predictions made by IPA software based off genes which were found to be dysregulated in epithelial cells of 6-month-old *Prickle1$^{f/f}$ MiCre* mice cKO (n=3) in diestrus.

**Supplemental Table 7. Pathway analysis of predicted diseases and disorders associated with dysregulated genes in the epithelial cell population (clusters 2 and 11) of *Prickle1$^{f/f}$* MiC cKO mice.**

| **Diseases and Disorders** | ***P* value** |
|---|---|
| Organismal injury and abnormalities | 1.34E-58 |
| Cancer | 3.59E-54 |
| Reproductive system disease | 3.59E-54 |
| Gastrointestinal disease | 1.09E-47 |
| Endocrine system disorders | 9.72E-43 |

Function predictions made by IPA software based off genes which were found to be dysregulated in epithelial cells of 6-month-old *Prickle1$^{f/f}$ MiCre* mice cKO (n=3) in diestrus.

**Supplemental Table 8. Pathway analysis of predicted molecular and cellular functions associated with dysregulated genes in the epithelial cell population (clusters 2 and 11) of *Prickle1$^{f/f}$* MiC cKO mice.**

| **Molecular and Cellular Functions** | ***P* value** |
|---|---|
| Cellular movement | 2.19E-67 |
| Cell death and survival | 2.43E-48 |
| Cellular development | 1.62E-35 |
| Cellular growth and proliferation | 1.62E-35 |
| Cell-to-cell signaling and interaction | 1.60E-27 |

Function predictions made by IPA software based off genes which were found to be dysregulated in epithelial cells of 6-month-old *Prickle1$^{f/f}$ MiCre* mice cKO (n=3) in diestrus.

**Supplemental Table 9. Pathway analysis of predicted physiological system development and functions associated with dysregulated genes in the epithelial cell population (clusters 2 and 11) of *Prickle1*$^{f/f}$ MiC cKO mice.**

| **Physiological System Development and Function** | ***P* value** |
|---|---|
| Organismal survival | 1.34E-58 |
| Cardiovascular system development and function | 2.15E-49 |
| Organismal development | 2.15E-49 |
| Tissue morphology | 4.00E-38 |
| Immune cell trafficking | 1.86E-37 |

Function predictions made by IPA software based off genes which were found to be dysregulated in epithelial cells of 6-month-old *Prickle1*$^{f/f}$ *MiCre* mice cKO (n=3) in diestrus.
